# Whole Genome Sequencing Reveals *Enterobacter hormaechei* as a Key Bloodstream Pathogen in Six Tertiary Care Hospitals in Southwestern Nigeria

**DOI:** 10.1101/2024.12.02.626058

**Authors:** Faith I. Oni, Ayorinde O. Afolayan, Anderson O. Oaikhena, Erkison Ewomazino Odih, Odion O. Ikhimiukor, Veronica O. Ogunleye, Aaron O. Aboderin, Olatunde F. Olabisi, Adewale A. Amupitan, Abayomi Fadeyi, Rasaki A. Raheem, Bashirat A. Olanipekun, Charles J. Elikwu, Oluwadamilola A. Sadare, Phillip O. Oshun, Oyinlola O. Oduyebo, Folashade Ojo, Abolaji T. Adeyemo, Ifeanyi E Mba, Abiodun Egwuenu, Tochi Okwor, Anthony Underwood, Silvia Argimón, Chikwe Ihekweazu, David M. Aanensen, Iruka N. Okeke

## Abstract

*Enterobacter* spp. are an important cause of healthcare-associated bloodstream infections uncommonly reported in Africa. This study used whole genome sequencing (WGS) to characterise *Enterobacter* spp. from hospitals in Nigeria’s antimicrobial resistance (AMR) surveillance system. Blood-culture isolates of *Enterobacter* from six such tertiary-care hospitals recovered between 2014 and 2020 were re-identified and antimicrobial susceptibility-tested using VITEK2. Illumina technology provided whole genome sequences for genome nomenclature, antimicrobial resistance gene prediction, Single Nucleotide Polymorphism (SNP) phylogeny, and multi-locus sequence typing via publicly available bioinformatics pipelines. Initial biochemical delineation often misclassified *Enterobacter*, necessitating whole-genome sequencing for accurate classification. Among 98 *Enterobacter* received, *Enterobacter hormaechei* subspecies *xiangfangensis* predominated (43), followed by other *E. hormachei* subspecies (18), *E. cloacae* (26), *E. roggenkampii* (4), *E. bugandensis* (3), *E. kobei* (2), *E. asburiae* (1) and *E. cancerogenous* (1). Cephalosporins, aminoglycoside, chloramphenicol, macrolide, and carbapenem resistance in *E. hormaechei* was attributed to known resistance genes. They belonged to clusters III, IV, and VIII based on *hsp60* typing and clades A, B, C, and D according to Sutton and Co’s nomenclature. These isolates and other *Enterobacter* species recently reported from Nigeria reveal extensive *E. hormaechei* diversity, as well as clusters representing potential outbreaks*. Enterobacter hormaechei*, often misidentified and rarely reported from Nigeria, is this study’s most common blood culture isolated *Enterobacter* spp. Uncovering underappreciated species as important bloodstream pathogens and retrospective detection of likely outbreaks emphasise the value of genomic surveillance in resource-limited settings.

**DATA SUMMARY:** All sequence reads were submitted to the European Nucleotide Archive (ENA) under the project ID PRJEB29739 (https://www.ebi.ac.uk/ena/browser/view/PRJEB29739). Accessions can be found in Table S1.

**IMPACT STATEMENT:** Accurate identification of *Enterobacter* is essential in healthcare settings as misidentification can lead to selecting antimicrobials to the genus is intrinsically resistant resistant before susceptibility testing results are available. Also, misidentification can compromise microbiology support for infection prevention and control. We show that *E. hormaechei*, which is never reported from clinical laboratories in Nigeria, is frequently misidentified using conventional methods like tube- or strip biochemical testing and VITEK systems. Whole genome sequence data demonstrates that *E. hormaechei* and *E. cloacae* are the most common *Enterobacter* isolated from bloodstream infections in Nigeria. Enhanced identification methods for surveillance play pivotal roles in improving patient care, optimising antibiotic stewardship, and combating the evolving challenges posed by this pathogen. Overall, this study reveals the effectiveness of WGS in correctly identifying this important pathogen.

## INTRODUCTION

The *Enterobacter* genus, comprising Gram-negative, non-spore-forming bacilli belonging to the *Enterobacteriaceae* family, are natural commensals of the gut of humans and animals [1] and are also commonly isolated from environmental sources such as water, sewage, soil, and plants [2]. The genus is a member of the ‘ESKAPE’ group of pathogens (*Enterococcus*, *Staphylococcus*, *Klebsiella*, *Acinetobacter*, *Pseudomonas,* and *Enterobacter* species) known to exhibit multidrug resistance [3] commonly and are an increasingly common cause of opportunistic nosocomial infections, but can also cause community-acquired infections [4]. *Enterobacter* can infect multiple sites, causing urinary tract, cerebral abscess, wounds, pneumonia, meningitis, abdominal and surgical site infections [5], as well as bloodstream infections [6,7].

An important sub-group among the *Enterobacter* species is the *Enterobacter cloacae* complex (ECC), comprising six species: *Enterobacter cloacae*, *Enterobacter asburiae*, *Enterobacter hormaechei*, *Enterobacter kobei*, *Enterobacter ludwigii* and *Enterobacter nimipressuralis* [6]. These species often evade precise identification due to the challenge of accurately biochemically profiling them using tube-based biochemicals commonly used in resource-limited settings [8]. Automated biochemical systems like the VITEK2 system and Matrix-Assisted Laser Desorption Ionization-Time of Flight mass spectrometry (MALDI-TOF) [6] also frequently misclassify *Enterobacter* species [9].

Whole genome sequencing (WGS) emerges as a promising solution for precise species-level identification, as demonstrated in recent studies ([9][10]. WGS enables a deeper understanding of ECC epidemiology, revealing cryptic species like *E. hormaechei* subsp. *xiangfangensis*, frequently misclassified as *E. cloacae* by traditional methods [11]. Notably, ECC is identified as a major Gram-negative bacterium responsible for neonatal sepsis in low- and medium-income countries like Nigeria [10]. However, many bloodstream-associated *Enterobacter* infections in Africa are reported with limited, if any, speciation or sub-speciation data [12], [13], [14]. In Nigerian hospitals, necessary reliance on biochemical tests, particularly tube-based biochemicals, often results in misidentification and inadequate species-level resolution for this important, AMR priority pathogen. Nigeria’s National AMR surveillance system was launched in 2017, and the Nigerian arm of the Global Health Research Unit (GHRU) for genomic surveillance of AMR provides WGS-based reference laboratory services to sentinel laboratories across the country[15], [16].

We used WGS to identify and characterise the prevalent *Enterobacter* species isolated from bloodstream infections in selected Nigerian hospitals between 2014 and 2020, and uncovered limitations of conventional biochemical methods.

## METHODS

### Collection of Presumptive bloodstream *Enterobacter* sp.

*Enterobacter* strains were isolated from blood cultures collected between 2014 and 2020 from six tertiary-care hospitals registered in the Nigerian Antimicrobial Resistance (AMR) surveillance system. Metadata containing preliminary information for each isolate was received from the contributing hospitals. From a total of 2,383 isolates processed by the reference laboratory, 63 were presumptively identified by the sentinels as various species of *Enterobacter* (*E. cloacae* (n=29), *E. aerogenes* (n=27), *E. gergoviae* (n=5), *E. agglomerans* (n=1) and *E. hormaechei* (n=1)). An additional 80 isolates not identified to species level were submitted as Enterobacteriaceae.

### Re-identification and antimicrobial susceptibility testing

Strains received through the antimicrobial resistance surveillance system were plated out on MacConkey agar to assess colony purity and phenotype. Heterogenous cultures were purified and strains were re-identified using the VITEK2 GN ID cards (21341). Their antimicrobial susceptibility profile was also determined using GN AST cards (N280 414531) that test for susceptibility to ampicillin, amikacin, gentamicin, cefuroxime, amoxicillin/clavulanic acid, cefepime, ceftriaxone, piperacillin/tazobactam, nitrofurantoin, cefuroxime_axetil, ciprofloxacin, nalidixic acid, meropenem, ertapenem, imipenem, tigecycline, trimethoprim-sulphamethoxazole, and colistin. The results were interpreted according to the CLSI [17].

### DNA extraction, Library preparation, and Whole Genome Sequencing

DNA of isolates was extracted using the Wizard DNA extraction kit (Promega, Wisconsin, USA) (A1125) according to the manufacturer’s protocols. Extracted DNA was quantified using the Qubit dsDNA BR Assay Kit (Invitrogen, Waltham, MA, United States). Libraries were prepared using the NEBNext Ultra II FS DNA library kit for Illumina with 384 unique indexes (New England Biolabs, Ipswich, MA, United States). Double-stranded DNA libraries were then sequenced using the HiSeq X10 with 150 bp paired-end chemistry (Illumina, San Diego, CA, United States).

### Whole Genome Sequence analysis

All sequence analyses were carried out using GHRU protocols (https://www.protocols.io/view/ghru-genomic-surveillance-of-antimicrobial-resista-bp2l6b11kgqe/v4). Genome assembly and quality control were carried out using the *de novo* assembly pipeline in the GHRU protocol. Assembly metrics were: N50 score, >50000; number of contigs that are >= 0 bp, <500; number of contigs that are >= 1000 bp, <300; Total length (>= 1000 bp), >4096846 or <6099522 and percentage_contamination <5.

Speciation and selection of reference for single nucleotide polymorphism phylogeny were done using the Bactinspector (check_species and closest_match) tool (https://gitlab.com/antunderwood/bactinspector). Pathogenwatch (https://pathogen.watch/ - v21.4.3 [18] was used to validate species identification. The closest reference genome selected for *E. hormaechei* was NZ_CP017183.1 (https://www.ncbi.nlm.nih.gov/nuccore/NZ_CP017183.1) while NZ_CP009756.1(https://www.ncbi.nlm.nih.gov/nuccore/NZ_CP009756.1) was chosen for *E. cloacae*. Mapping to reference was done with the bwa mem tool (https://arxiv.org/abs/1303.3997). Variant calling and filtering were done with samtools/bcf tools (https://github.com/samtools/bcftools), and maximum likelihood phylogenetic trees were constructed. The pair-wise SNP distances for likely outbreak isolates were calculated using FastaDist (https://gitlab.com/antunderwood/fastadist).

Identification of multilocus sequence types (according to the Pasteur scheme) was done using the ARIBA software [19] and the PubMed database (https://www.protocols.io/view/ghru-genomic-surveillance-of-antimicrobial-resista-bpn6mmhe).

Antimicrobial resistance genes, virulence genes, and plasmid replicons were predicted *in silico* using the aforementioned GHRU protocol. Predicted genes tagged as “yes” or “yes_nonunique” by the ARIBA software were accepted as present in the genomes. The criteria used for defining Multidrug resistance (MDR) in isolates according to [20]: non-susceptibility to ≥1 agent in >3 antimicrobial categories [20]. AMRFinderPlus version 3.1.0 [21] and the Comprehensive Antimicrobial Resistance Database (CARD) [22] were used to determine *ampC* variants among *Enterobacter spp*.

Hoffman clustering of *Enterobacter spp*. was done using hsp60ECC tool (https://github.com/karubiotools/hsp60ECCtool). Publicly available data from the Sands *et al.* 2021 study were retrieved from the European Nucleotide Archive (ENA) under the project accession number SAMEA7472464. Fastq files of *Enterobacter* spp. isolated from Nigeria were downloaded from ENA (https://www.ebi.ac.uk/ena/browser/view/SAMEA7472464?show=reads) and assembled using the aforementioned GHRU de novo assembly protocol. Hoffman clustering and clades were determined for the species using the HSP60ECC tool.

The average Nucleotide Identity (ANI) of genomes was obtained using the FastANI tool (https://github.com/ParBLiSS/FastANI) [23]. The “many-to-many” method in FastANI was used to compute ANI between multiple query genomes (genomes from this study) and multiple reference genomes (Sands *et al.* Enterobacter Nigeria genomes)[24].

Novel STs identified in this study were first confirmed as novel by querying their fasta sequences on the PubMLST Public database for molecular typing and microbial genome diversity (https://pubmlst.org/bigsdb?db=pubmlst_ecloacae_seqdef). Their profiles were then submitted, and STs were assigned as follows: G20500026: assigned - ST-1995, G20500682: assigned - ST-1996, G18503215: assigned - ST-1997, G18503415: assigned - ST-1998 and G18503407: assigned - ST-1998.

For data visualisation, iTOL (https://itol.embl.de/tree/197211635839451656920611#) [25], itol.toolkit R package (https://github.com/TongZhou2017/itol.toolkit) [26] and Microsoft Excel version 16.62 (2022) were used.

For map drawing, R packages—naijR (https://docs.ropensci.org/naijR/articles/nigeria-maps.html), SF (https://cran.r-project.org/web/packages/sf/index.html), map (https://cran.r-project.org/web/packages/tmap/index.html) and feathers (https://cran.r-project.org/web/packages/feather/index.html) were used.

## RESULTS

### Enterobacter species identified by WGS

Sentinel labs sent a total of 63 isolates as *Enterobacter* spp., and of these, 27 were verified as *Enterobacter* by VITEK2, out of which WGS eventually identified 13 as belonging to the genus. 85 more isolates were sent as ‘Enterobacteriaceae’, species belonging to other families, or unidentified, and identified by WGS as *Enterobacter* (Figure 1c). In total, therefore, 98 (4.57%) *Enterobacter* isolates from between the years 2014 and 2019 were received at the national reference laboratory as *Enterobacter*, which represented the sixth most common bloodstream-isolated genus (after *Klebsiella, Escherichia, Staphylococcus, Acinetobacter,* and *Salmonella*) (Figure 1a). Of these 98 isolates, 61 (62.25%) were identified by WGS as *E. hormaechei*, 26 (26.53%) as *E. cloacae*, and the rest, 11 (11.22%) were identified as *E. roggenkampii* (4), *E. bugandensis* (3), *E. kobei* (2), *E. asburiae* (1) and *E. cancerogenous* (1) (Figure 1b).

**Figure 1a:**
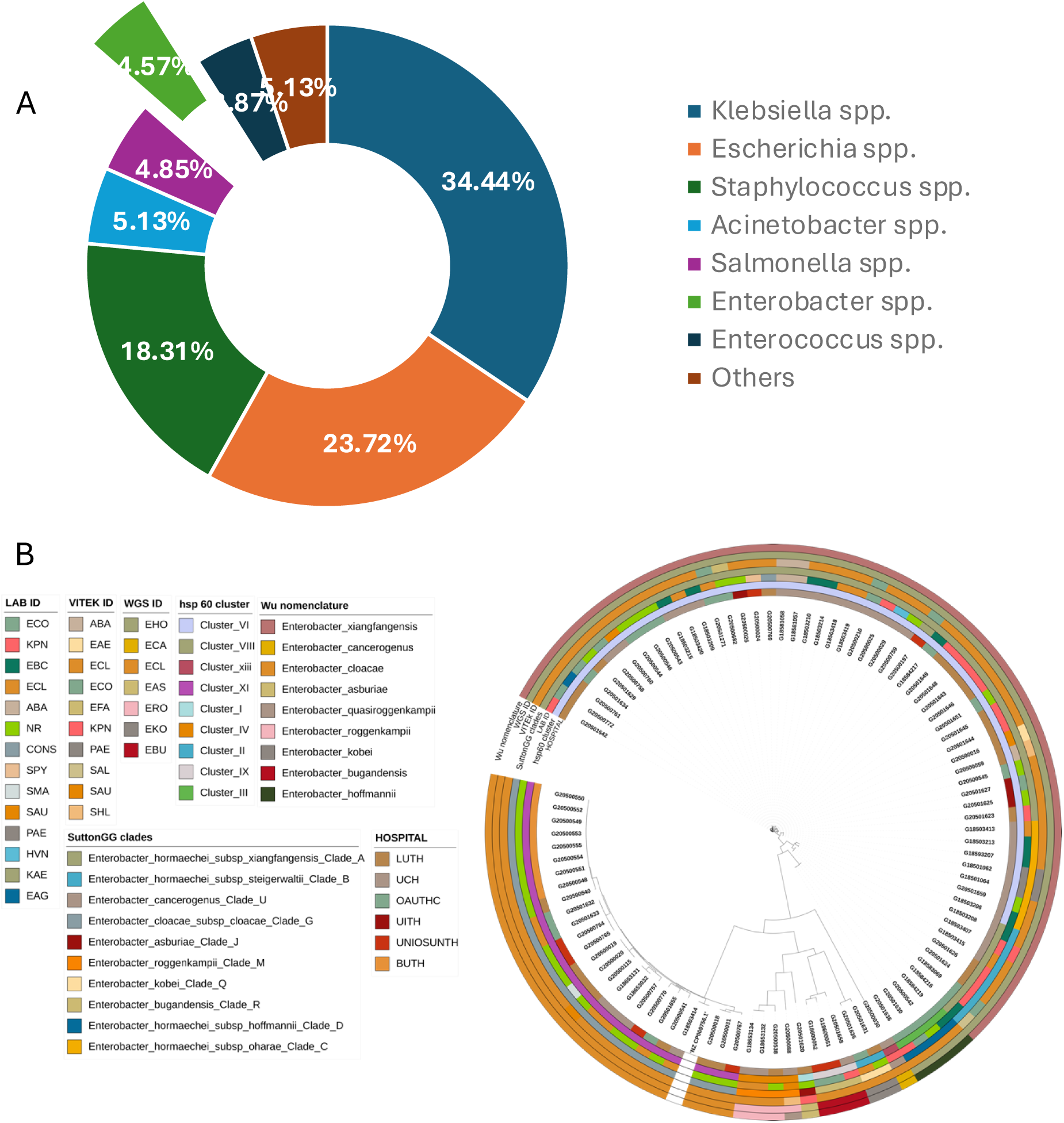
Proportion of Genera received from invasive infections from six sentinels in the Nigeria AMR surveillance system (2014-2020) Figure 1b: Maximum likelihood tree showing whole genome sequence identifications of *Enterobacter* isolates recovered from patients admitted into six hospitals in southwestern Nigeria juxtaposed on identifications of *Enterobacter* species by the diagnostic lab sentinels and reference lab Vitek2 reidentification, their clades and Hoffman clusters.

Forty-nine of the 61 WGS-identified *E. hormaechei* isolates were identified as *Klebsiella pneumoniae* (n=14, 29%), Enterobacteriaceae (n=11, 22%), *Enterobacter cloacae* complex (n=6, 12%), *Escherichia coli* (n=6, 2%), *Acinetobacter baumannii* (n=3, 6%), *Staphylococcus aureus* (n=2, 4%), *Pseudomonas aeruginosa* (n=3, 6%), *Pantoea agglomerans* (n=1, 2%), Coagulase negative *Staphylococcus* (n=1, 2%), *Streptococcus pyogenes* (n=1, 2%) and *Halovenus* (n=1, 2%) at the sentinel laboratories (Figure 2a).

**Figure 2:**
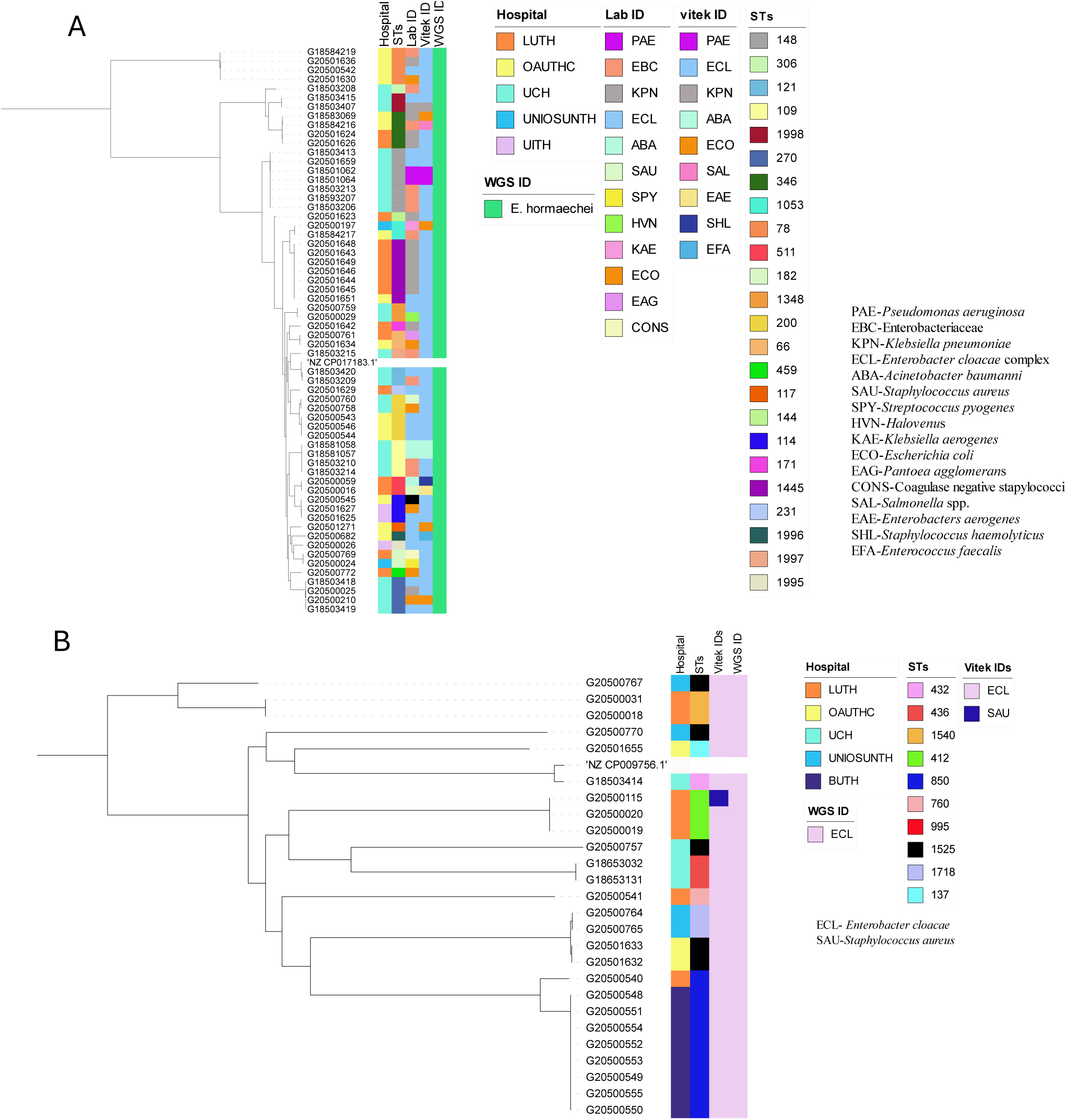
Initial Identification of the two most frequently-detected *Enterobacter* species by Reference and Sentinel Laboratories, whole genome sequence ID and their sequence types (a) *Enterobacter hormaechei* (b) *E. cloacae*

At the national reference laboratory, the 61 WGS-identified *E. hormaechei* were initially identified (using VITEK2) as *Enterobacter cloacae* complex (n=48, 79%), *Escherichia coli* (n=4, 7%), *Acinetobacter baumannii* (n=2, 3.3%), *Enterobacter aerogenes* (n=1, 1.6%), *Salmonella* spp. (n=1, 1.6%), *Pseudomonas aeruginosa* (n=1, 1.6%) and *Klebsiella pneumoniae* (n=1, 1.6%). Selection of Gram-positive VITEK cards after Gram miscalling meant that three further isolates were misidentified as Enterococci or Staphylococci: *Enterococcus faecalis* (n=1), 1.6%, *Staphylococcus haemolyticus* (n=1, 1.6%) and *Staphylococcus aureus* (n=1, 1.6%), (Figure 2a, Table S2). VITEK2 percentage probabilities of identification of the species are shown in Table S2. The proportion of *E. cloacae* (the second most abundant species identified among the *Enterobacter* spp.) identified biochemically, using VITEK2 as *E. cloacae* was 96.1% (n=25) while 3.9% (n=1) was identified as *Staphylococcus aureus*. The sensitivity, specificity, positive predictive and negative predictive values (100%, 0%, 100%, and 0% respectively) [27, 28] for *E. cloacae* complex identification in this study by VITEK2 using GN ID cards (21341), show that this method is adequate for *E. cloacae* but suboptimal for *E. hormachei* (0% for all four values) in our setting (table S6).

Altogether, retrospective (2017 and prior) and prospective *Enterobacter* isolates were received from six tertiary hospitals: University College Hospital, Ibadan-UCH (n=35), Lagos University Teaching Hospital, Lagos-LUTH (n=30), Obafemi Awolowo University Teaching Hospital, Ile-Ife- OAUTHC (n=18), Babcock University Teaching Hospital, Ogun-BUTH (n=8), Osun State University Teaching Hospital, Osogbo- UNIOSUNTH (n=4) and University of Ilorin Teaching Hospital, Ilorin- UITH (n=3) (Figure 1b, 1c).

*Enterobacter hormaechei* strains—the most abundant *Enterobacter* species identified—were detected in 5 of the six sentinels and were quite diverse (Simpson’s diversity index = 0.937) as they belonged to 22 different sequence types (STs) (Figure 2a), including the novel STs, ST1995 (n=1), ST1996 (n=1), ST1997 (n=1), and ST1998 (n=2). This species was most commonly identified from UCH, representing 25/35 of the *Enterobacter* species (Figure 2a) and encompassing 7 STs, including two novel ST- ST1997 and ST1998. *E. hormaechei* were also retrieved from the sentinel sites in Ile-Ife (OAUTHC; 15 strains belonging to 9 STs including a novel ST- ST1996), Lagos (LUTH; 16 strains belonging to 9 STs), Ilorin (UITH; 3 strains belonging to STs 114 and novel ST1995), and Osogbo (UNIOSUNTH; 2 strains belonging to STs 1053 and 182) (Figure 2a). No particular STs were seen to be shared across the hospitals. The STs 148 and 1445 were the most prevalent among the isolates.

*Enterobacter cloacae* (n=26) were collected from UCH (n=4), LUTH (n=7), UNIOSUNTH (n=4), BUTH (n=8), and OAUTHC (n=3). Isolates belonged to 10 different STs- 432 (1), 436 (1), 1540 (2), 412 (3), 850 (9), 760 (1), 995 (1), 1525 (5), 1718 (2), and 137 (1) with ST850 being most prevalent. 3 STs (432, 436, and 995) were detected among strains from UCH, 4 (1540, 412,760 and 850) in LUTH, 1 (1718) in UNIOSUNTH, 1 (850) in BUTH, and 2 (137 and 1525) in OAUTHC. *E. cloacae* ST850 was commonly found in BUTH and LUTH (Figure 2b).

#### Antimicrobial resistance genes and phylogenetic relationships among *Enterobacter* isolates

Carbapenems, beta-lactams, fluoroquinolones, cephalosporins, aminoglycosides, and colistin are antibiotics commonly used to treat infections caused by Enterobacter species. [29], [30]. Genes responsible for resistance to these antibiotics - carbapenems (*bla*_NDM_ - 5), beta-lactams/ cephalosporins (*bla*_ACT-45_ - 61, *bla*_TEM,_ - 39, *bla*_CTX-M-15_ - 33, *bla*_OXA_ - 40, *bla_SHV_* - 3, *bla_DHA_* - 2), fluoroquinolones (*aac(6’)-Ib-cr* - 38 *, qnrB1* - 32), fosfomycin (*fosA* - 49), chloramphenicol (*catA1* - 29*, catA2* - 9*, catB3* - 8), macrolide (*mphA* - 9*, mphE* - 4*, msrE* - 4) sulfonamide (*sul1* - 19, *sul2* - 38), tetracycline (*tet(38)* - 1*, tet(A)* - 38*, tet(D)* - 1*, tet(K)* - 1), quinolone (*qnrB1*- 32, *qnrB4* - 2, *qnrS1* −13), trimethoprim (*dfrA1* - 4, *dfrA12* - 5, *dfrA14* - 44, *dfrA15* - 1, dfrA27 - 1*, dfrG* - 1), quartenary ammonium compounds (*qacEdelta1* - 18), aminoglycosides (*aadA1* - 25*, aadA2* - 5*, aph(3’)-Ia* - 2 *, armA* - 4 *, aac (3)-Ile* - 29 *, aph(3”)- Ib* - 40, *aph(6)-Id* - 38), and colistin (mcr10.1 - 2) - were detected in *hormaechei* genomes (n=61) (Figure 3a). Fifty-six strains were classified as multidrug-resistant (as defined by Magiorakos *et al.,* 2012[20]) due to the *in-silico* detection of more than two of these genes. *Enterobacter* spp. are known to carry core chromosomal AmpC-type beta-lactamases and their variants [31] (Table S4). While various other *bla*_ACT_ alleles have been associated with *E. hormaechei* in the literature [31][32], by using ResFinder to analyse the resistance genes, we found all 61 isolates in this study carried *bla*_ACT-45_. This gene was earlier reported to occur naturally in *E. hormaechei* subsp. *xiangfangensis* and contribute to antibiotic resistance mechanisms observed in these strains [33]. Meanwhile, further analysis of the *ampC* variants using AMRfinder plus and CARD revealed 27 different *ampC* variants in the 98 isolates in this study (Table S4). The most common variant in this study was bla_ACT-16_ and was found to be associated with *E. hormaechei*. Of the 98 *Enterobacter* strains we sequenced, all but 16 isolates carried, in addition to core AmpC-type beta-lactamases, one or more acquired beta-lactamase genes associated with mobile elements as well as other resistance genes that confer resistance to the aminoglycosides, trimethoprim, colistin, fluoroquinolone, fosfomycin, chloramphenicol, macrolide, sulfonamide, and tetracycline (Figure 3a). Thirty-three of the *E. hormachei* and 18 *E. cloacae* carried *bla*_CTX-M-15_. In 12/51 of these cases, an IncFIb plasmid replicon was also detected. IncFIA_HI1, IncHI2, IncHI2A, and IncR plasmid replicons were also detected among the *bla_CTX-M-15_* carrying strains. Carbapenemase gene *bla*_NDM-1_ was found in five of the *E. hormaechei* genomes. *dfrA* alleles conferring trimethoprim resistance were almost ubiquitous, with *dfrA14* predominating in 51/61 and 22/26 *E. hormachei* and *E. cloacae* genomes, respectively.

**Figure 3:**
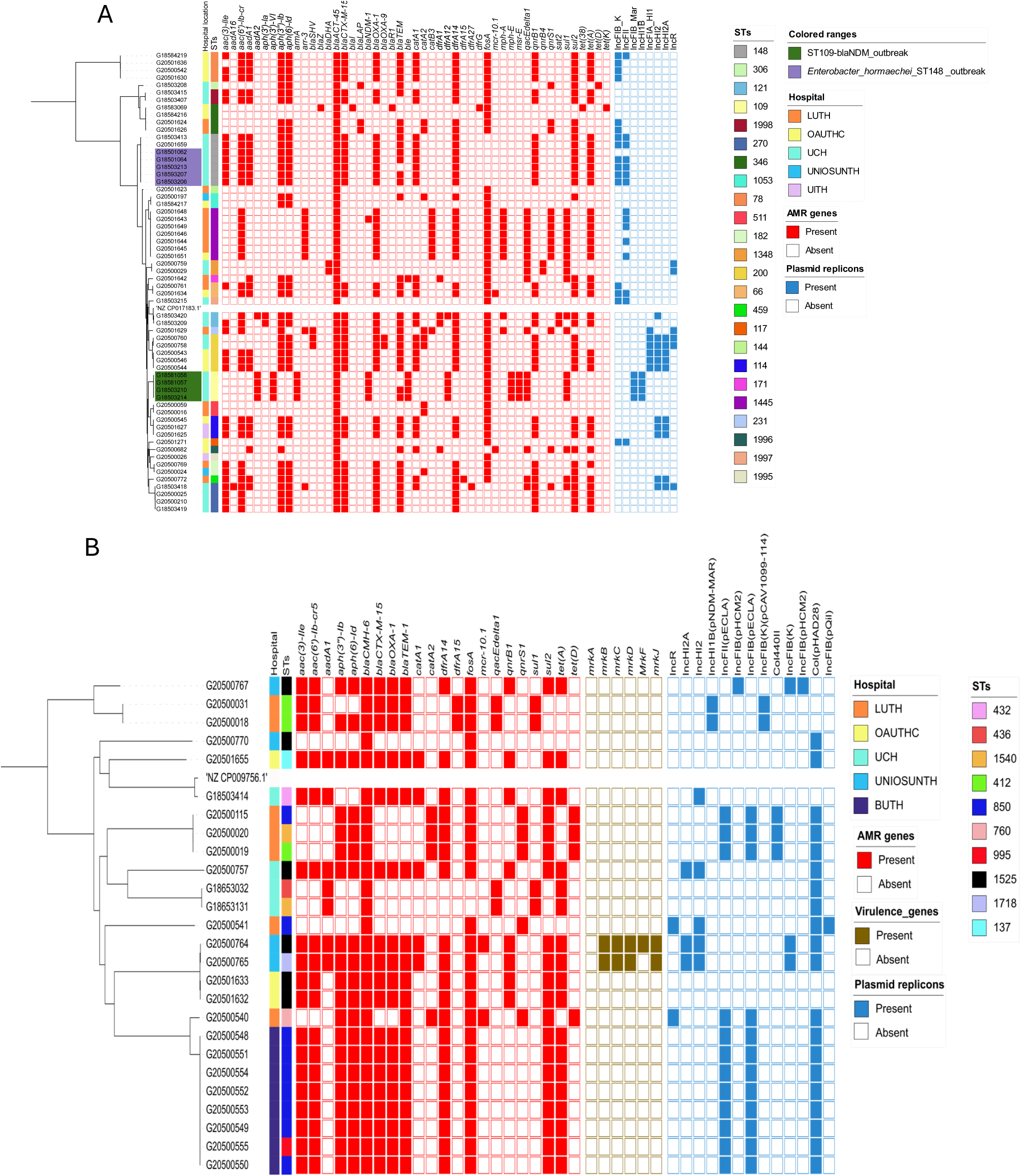
Maximum likelihood phylogenetic trees showing the relationship among *Enterobacter* isolates belonging to the most frequently encountered species from different hospitals, sequence types, AMR genes, and plasmid replicons detected. (a) *Enterobacter hormaechei* (b) *E. cloacae*

The mobile colistin resistance *mcr*10.1 gene was found in two *hormaechei* strains. The *mcr10.1*-carrying *hormaechei* strains belonged to ST 66 and novel ST 1996 and had no plasmid replicon type in common, according to the output from plasmid finder. The phylogeny (Figure 3a) shows that isolates from the same location cluster together, often carrying identical resistance genes, suggesting very local epidemiologies for *E. hormaechei* lineages and that the predominance of this species throughout the whole network is not due to clonal expansion of one or a few clones.

We observed that *E. cloacae* strains (Figure 3b) were multidrug-resistant. Frequencies of resistance genes found in isolates (n=26) for aminoglycoside *(aac(3)-Ile* - 18*, aac(6’)-Ib-cr5* - 18*, aadA1* - 8*, aph(3”)-Ib* - 20*, aph(6)-Id* - 20), beta-lactams (*bla_CMH-6_* - 26*, bla_CTX-M-15_* - 18*, bla_OXA-1_* - 18*, bla_TEM-1_* - 19), chloramphenicol (*catA1* - 5*, catA2* - 4), sulphonamides (*sul1* - 4*, sul2* - 20), tetracycline (*tetA* - 18*, tetD* - 4), trimethoprim (*dfrA14* - 20*, dfrA15* - 2), Fosfomycin (*fosA* - 24), quinolone (*qnrB1* - 8*, qnrS1* - 5), and colistin (*mcr-10.1* - 2). Mobile colistin resistance gene *mcr10.1* was observed in two *E. cloacae* isolates belonging to ST1718 and ST850 isolated from UNIOSUNTH and LUTH, respectively (Figure 3b). These strains had otherwise different resistance gene and plasmid replicon profiles.

We calculated the concordance between the phenotypic AMR (Vitek2 AMR result) and the genotypic AMR (WGS AMR result) for the *E. cloacae* and *E. hormaechei* species (Table S3). A concordance of 1 signifies a 100% agreement between phenotypic and genotypic antibiotic resistance. For *E. hormaechei* resistance data, ampicillin, cefuroxime, cefuroxime axetil, Imipenem, meropenem, and amikacin showed 100% concordance results. On the other hand, Cefepime, cefoperazone/ sulbactam showed less than 50% concordance, while the other antibiotics showed greater than 50% but less than 100% concordance results. For *E. cloacae*, piperacillin/ tazobactam, cefoperazone/ sulbactam, amikacin, and colistin showed poor concordance and reflect that care needs to be taken in the choice of beta-lactam antimicrobials used for phenotypic testing in particular.

#### Potential *Enterobacter* health-care associated infection outbreaks detected

Two potential hospital outbreaks were retrospectively detected in this study among the *E. hormaechei* strains at the UCH facility. An ST109 cluster carrying the *bla*_NDM-1_ carbapenemase gene (Figure 3a) was comprised of strains that were phenotypically sensitive to meropenem, imipenem, and ertapenem with MICs of <= 0.25, 1 & 0.5 and <= 0.5 respectively (Table S5). These isolates also did not demonstrate phenotypic resistance to other beta-lactams attributable to *bla*_ACT-45_. The SNP distance among ST148-bearing strains was between 0 and 1 while the range of SNP distance between these putative outbreak isolates and *E. hormaechei* that are not part of the outbreak is between 144 - 258 SNPs (Figure S1). The ST109 putative outbreak strains carried aminoglycoside-resistance genes not seen in any of the other Enterobacter hormachei - *aph(3’)-VI* and *armA* as well as *aadA2*, *aph(3’)-Ia*, *bla*_ACT_, *bla*_NDM_, *bla*_TEM_, *ble*, *dfrA12*, *fosA*, *mphE*, *msrE* and *sul1* genes. Only one isolate outside this likely outbreak cluster carried *bla*_NDM-1_. It belonged to ST1445, was from a different facility, and contained a completely different repertoire of resistance genes (Figure 3a). The outbreak strains carried resistance genes belonging to six antimicrobial classes - aminoglycoside, beta-lactamase, trimethoprim, carbapenem, macrolide, and sulfonamide resistance -compared to the median number of resistance classes conferred by genes in non- ST109 strains (range 3-5). All four genomes contained IncFIB_Mar and IncHIB plasmid replicons not seen in other isolates.

A second cluster of 5 ST148 strains, also from UCH, harboured IncFIB_K and IncFII plasmid replicons and carried genes conferring resistance to aminoglycosides (*aac(6’)-Ib-cr, aac (3)-Ile*, *aadA1*, *aph(3”)-Ib*, *aph(6)-Id*), beta-lactams (*bla*_ACT45_, *bla*_CTX-M-15_, *bla*_OXA-1_, *bla*_TEM_), phenicol (*catA1*) quinolones (*qnrB1*), sulfonamides (*sul2*), trimethoprim (*dfrA14*) and tetracyclines (*tet*A) (Figure 3a). SNP distance among strains within the ST148 cluster was between 0 and 1 while the range of SNP distance between these outbreak isolates and the other strains is between 27542 and 31513 (Figure S1). They were phenotypically resistant to cephalosporin, carbapenem, aminoglycoside, quinolone, and trimethoprim (Table S5).

Sentinel laboratories in Nigeria can request accelerated sequencing of suspected outbreak clusters. [16]. However, although these likely outbreaks, for which retrospective time (other than year) and place information are not available, occurred at the sentinel that had the greatest success at identifying *Enterobacter* genus strains, both clusters contained isolates that were misclassified as different species at the sentinel and reference lab (VITEK2) levels, which would have hampered WGS-independent cluster identification.

#### Genomic context of Enterobacter genomes from this study and *Enterobacter* genomes from Sands *et al*., 2021 study

To enable us to analyse the distribution of *Enterobacter* lineages nationally, we downloaded all *Enterobacter* genomes associated with Nigeria that can be retrieved from the European Nucleotide Archive under project number PRJEB33565. All the genomes not from the current study (see Table S1) arose from the study of Sands *et al.,* a rigorous WGS-based neonatal sepsis study, and were submitted as *Enterobacter* species (19) – *cloacae* (17), *hormaechei subsp. xiangfangensis* (1) and *hormaechei* (1). These were included them in our ANI analysis. Four Sands *et al.* (2021) *E. cloacae* genomes clustered with *E. hormaechei* from this study, 3 with *E. roggenkampii* from this study, 1 with *E. bugandensis* from this study, and the others with *E. cloacae* from this study. One of Sands *et al*., 2021 *E. hormaechei* genomes clustered with E. *roggenkampii* and the other with E. *hormaechei* from this study (Figure 4a). We re-identified the Sands *Enterobacter* species using our assembly and speciation pipelines. Sands *et al. E. cloacae* were identified as *roggenkampii* (3), *hormaechei* (4), *bugandensis* (1), *cloacae* (9); the *E. hormaechei* subsp. Xiangfangensis as *E. hormaechei* and the *E. hormaechei* as *E. roggenkampii*. Sands *et al*., 2021 used both BLAST and Pathogenwatch to identify their bacteria species. The identities from Pathogenwatch of Sands *et al*., 2021 genomes and the genomes from our study correlated with the identities from our pipelines. The output from Hoffman’s classification yielded identities aligned with our pipeline’s results (Figure 4b).

**Figure 4.**
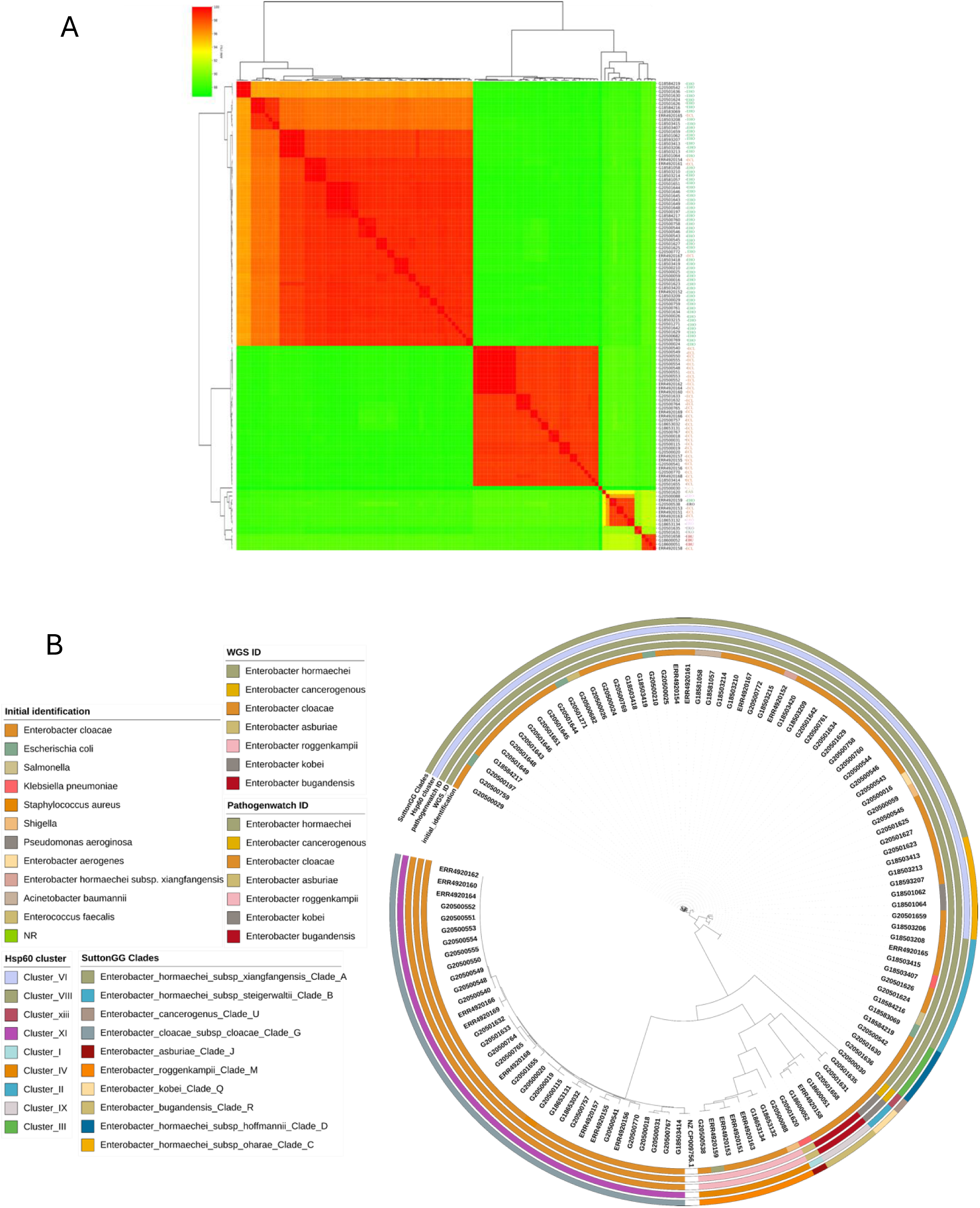
Comparison of genomes from this study and those of isolates from elsewhere in Nigeria. Genomes IDs are prefixed by ERR* if submitted by Sands *et al*. (2021) *or* G*(This study) (a): ANI analysis of genome-sequenced *Enterobacter spp.- E. hormaechei* (EHO), *E. cloacae* (ECL), *E. bugandensis* (EBU), *E. asburiae* (EAS), *E. roggenkampii* (ERO), *E. kobei* (EKO). (b) Maximum likelihood tree illustrating the phylogeny among all available Nigeria genomes in the European Nucleotide Archive

The Sands *et al*., 2021 [24] isolates were from northern Nigeria (National Hospital Abuja, Abuja (NN), Wuse District Hospital, Abuja (NW) and Murtala Muhammad Specialist Hospital, Kano (NK) in Nigeria), geographically distinct from the area where our isolates were collected. Altogether, the two studies identified 43 Enterobacter STs, 11 of which were found at more than one facility, and 2 (STs 109 and 850) were reported in this study and the Sands et al. study. Our data contain no clinical or outcome information on the isolates. However, Sands *et al.* found that the STs 1238, 850, 1031, and 544 (see their supplemental data Fig 9) were commonly associated with fatal infections. This study recovered isolates belonging to these STs from neonatal sepsis infections. The identities from all nine hospitals are shown on the Nigerian map (Figure 5).

**Figure 5:**
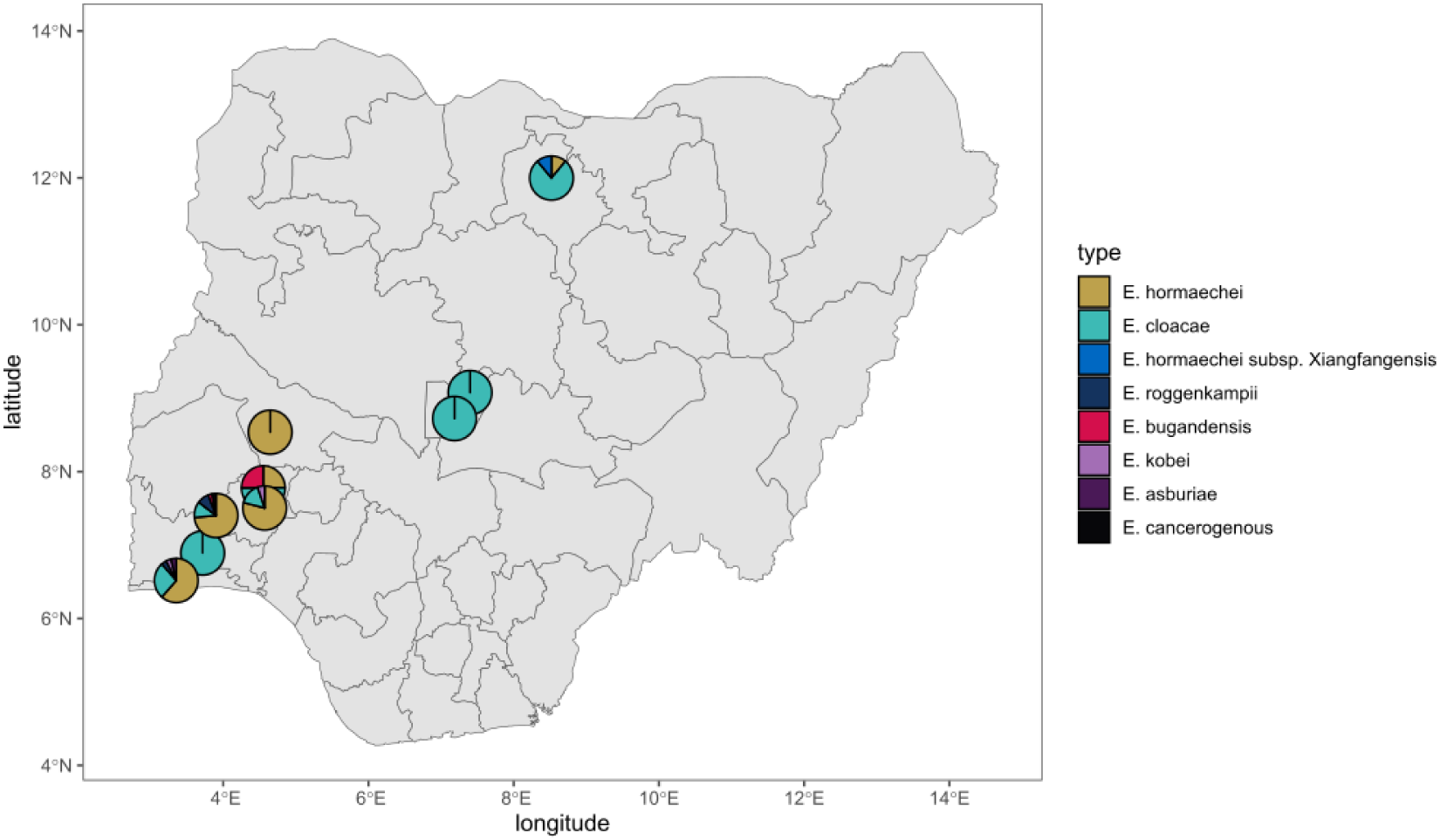
Geographic source of genome-sequenced *Enterobacter* in Nigeria. Isolates originated from southwestern Nigeria (This study) and north and central Nigeria (Sands et al, 2021)

## DISCUSSION

In this study, we performed whole genome sequencing of bloodstream isolates submitted to the Nigerian surveillance system. Bloodstream isolates collected between 2014 and 2020 included *Staphylococcus aureus, Escherichia coli, Klebsiella pneumoniae, Acinetobacter baumanii, Salmonella, Pseudomonas aeruginosa, Enterobacter spp.* as the most common genera (Figure 1a). Of 2360 isolates received, 63 were initially sent as presumptive *Enterobacter,* and 98 (4.2%) were eventually identified as *Enterobacter* species. Currently, there are 45 (22 named and 21 without assigned names) species of *Enterobacter* [5], [31], [34], seven were identified in this study (Figure 1b). Among the 22 named *Enterobacter* spp., 7 belong to the *Enterobacter cloacae* complex, and they include *E. cloacae, E. hormaechei, E. mori, E. asburiae, E. ludwigii, E. nimipressuralis* and *E. kobei*. All but *E. mori, ludwigii, and nimipressuralis* were identified in this study. We also identified *E. roggenkampii*, *bugandensis*, and *cancerogenous*, which are not members of the *cloacae* complex (Figure 1a).

*Enterobacter hormaechei* is frequently encountered in clinical specimens and is commonly considered a nosocomial pathogen [34] but there are only a few reports of bloodstream *Enterobacter hormaechei* infections from Africa. Duru *et al*., 2020 [12] reported the identification of *Enterobacter* spp. from blood samples, but the taxonomic resolution was limited to the genus level. In contrast to our study, which identified *E. hormaechei* as the most common *Enterobacter* species, a previous study conducted more than a decade ago [14] in Benin City, Nigeria, identified *E. sakazakii* and *E. aerogenes* as the most prevalent Enterobacter species from clinical samples, which included blood and did not report *Enterobacter hormaechei*. In the Sands *et al*.(2021) study, *Enterobacter* were prominent in Nigeria. The conspicuous dearth of any previous report on *E. hormaechei* from other clinical samples in Nigeria is likely due to the misidentification of *Enterobacter* species using biochemical identification methods.

Identifying the *Enterobacter* genus is often challenging [34]. The genus is often inaccurately identified by clinical laboratories using biochemical and other phenotype-based tests, VITEK2, and MALDI-TOF mass spectrometry [6, 35]. Resource-limited settings may face heightened challenges with identifying bacterial pathogens and therefore supportive reference laboratory services are critical [8, 16, 36]. In this study, four *Enterobacter* species were submitted as Gram-positive strains and three of those 4 belonged to the most common species- *Enterobacter hormaechei.* Isolates were submitted after identification by tube- or strip biochemical test-using labs as *Enterobacteriaceae* (n=11) or *Enterobacter cloacae* (n=5) or were misidentified by sentinels as *Acinetobacter baumannii* (n=3), *Pseudomonas* (n=2), *Klebsiella pneumoniae* (n=14), *Escherichia coli* (n=6), *Halovenus* (n=1), *Klebsiella aerogenes* (n=2)). At the reference laboratory, VITEK2 lacked the resolution to delineate these species as *Enterobacter hormaechei,* although it did place most of them in the *cloacae* complex. Only WGS identified these organisms accurately at the species and sub-species level. We show that *E. hormaechei* was commonly isolated in all the participating in this study’s hospitals (except BUTH from which overall, only a few isolates were obtained) (Figure 1b, 1c). The within-species misclassifications have little consequence for patient management but prevent early identification of clusters, which is important for infection prevention and control. Our results, and the important sub-specific nuances we found, emphasize the need to integrate WGS into routine clinical diagnosis of infectious diseases. Also, there is a need to update the VITEK2 and MALDI-TOF databases to improve the accuracy of speciation.

Analysis of the *hsp60* gene (a housekeeping gene) has conventionally been used to sub-classify the *Enterobacter cloacae* complex into 13 genetic clusters (Hoffman clusters I - XII and an unstable sequence crowd xiii) [37]. A whole genome analysis study (1,997 *Enterobacter* genomes) updated the taxonomy of the *Enterobacter* genus [11]. ANI thresholds between 94 and 96.5% and 97-98% for subspecies have good correlations with current species designations [38]. Average Nucleotide Identity (ANI) was used for classifying strains in the *Enterobacter cloacae* complex into 22 clades (A-V) which correspond to Hoffman clusters (I-XII) [39] (Figure 1b; Figure 4b). Making connections across the *Enterobacter* literature is challenging because of the different schemes, to which multi-locus sequence typing, offering much finer sub-classification, has been added. WGS approaches make it possible to easily classify strains according to all schemes, and therefore be able to compare disparate datasets. This made it possible for us to collate information from the Sands et al study, which was performed in different locations in Nigeria on an overlapping timescale. Like Sands et al [24], we found MLST most helpful.

Our isolates belonged to 22 different sequence types including novel sequence type (now assigned ST1995, ST1996, ST1997, and ST1998), which, with the 12 STs reported by Sands et al (of which 2 STs were seen in both studies) shows that the *E. hormaechei* population in Nigerian hospitals is considerably diverse. *E. hormaechei* strains were multidrug resistant with the detection of aminoglycoside, cephalosporin, chloramphenicol, macrolide, colistin, and carbapenem resistance genes. An ST148 and ST109-*bla*_NDM_ (Figure 3a) appear to represent outbreaks, comprised of isolates with SNP distances of ≤1 and ≤1 respectively. ST148 *Enterobacter* strains have been known to cause outbreaks in the past and have been identified among species isolated at hospitals. A study in Canada that investigated Carbapenemase-Producing Enterobacterales transmission clusters at a hospital system identified *bla*_VIM-1_-positive ST148 strains harbouring plasmid replicons IncR, HI2, and HI2A [40]. The ST148 strains in this study did not carry any carbapenemase gene but were multidrug-resistant (Figure 3a) and were phenotypically resistant to imipenem and meropenem with MICs of ≥8 and ≥16 (Table S5). This phenotype may be a result of overexpression of the chromosomal *ampC* gene alongside alteration in outer membrane transcriptome balance which is known to proffer other phenotypes such as carbapenem resistance [41]. OXA-48-like-producing ST109 *E. cloacae* was implicated alongside 22 *Klebsiella pneumoniae* and 3 *Escherichia coli* in outbreaks of OXA-48-like-producing *Enterobacteriaceae* in Czech hospitals in 2015. The ST109 *E. cloacae* strain harbored, in addition to the *bla*_OXA-48_ gene, *bla*_CTX-M-15_, *bla*_OXA-1_, and *bla*_TEM-1_ [42]. The ST109 strains in this study carried only the core *bla*_ACT-45_ and *bla*_NDM-1_ beta-lactamase genes (Figure 3). *E. cloacae* was identified in all the hospitals excluding UITH. A total of 26 isolates belonged to 10 different sequence types with the ST850 being most prevalent with SNP differences of ≤2. Nigeria was the only country from which Sands *et al.,*2021 recovered *Enterobacter* spp. from every sentinel, similar to our study. The ST859 *E. cloacae* strains from our study are not very distantly related to the ST850 *E. cloacae* genomes from Sands *et al*., with an SNP distance range of between 70 and 89. Altogether, they identified 3 ST850 *Enterobacter cloacae* among antimicrobial-resistant Gram-negative bacteria that cause neonatal sepsis in seven low- and middle-income countries, and all were reported from Nigeria. Unlike our study, Sands *et al* sentinels were in northern Nigeria. Thus, while our ST850 isolates appear to be focused at one sentinel, this clade may be circulating widely in Nigeria. *E. cloacae* isolates in this study were resistant to aminoglycosides, cephalosporins, and colistin. The presence of the mobile colistin resistance gene- *mcr-10.1* in two *E. cloacae* strains in this study is very worrisome as colistin, which is difficult to access in Nigeria, is one of the last available antibiotics used in the treatment of carbapenem-resistant infections and *mcr* genes are easily transmitted. Although there are few reports of colistin-resistant *Enterobacter* in Africa, colistin-resistant *E. cloacae* was recently identified in Sierra Leone. The strain belonged to ST850 and was resistant to cefazolin, gentamicin, and trimethoprim [43]. ST850 strains from this study were colistin-sensitive, and the colistin-resistant *E. cloacae* strains belonged to ST1525 and ST760. They also possessed *bla*_CTX-M-15_*, bla*_TEM1_*, bla*_CMH,_ *qnrS*, *qnrB*, *tetD*, *aph(6)-id*, *aadA1*, *dfrA*, *catA2*, *sul2* genes which confer resistance to beta-lactams, quinolone, tetracycline, trimethoprim, chloramphenicol, aminoglycoside, and sulfonamides.

The importance of *Enterobacter* species as bloodstream pathogens in Nigeria has heretofore been overlooked because precise identification of this genus is a difficulty for clinical laboratories due to limited biochemical capacity and the complicated taxonomy of the Genus. In this study, WGS allowed for the correct delineation of members of this genus, which uncovered *Enterobacter hormaechei* as the predominant species. It also allowed for retrospective identification of earlier missed outbreaks. In this study, the retrospective identification of potential outbreaks and the detection of genes conferring resistance to last-line drugs, carbapenems, and colistin is worrying.

The detection of *E. hormaechei* as the most prevalent *Enterobacter* bloodstream isolates and identification of key *E. cloacae* lineages in this study uncovers the need to strengthen clinical laboratory identification and for continued surveillance of this genus. This study emphasizes the importance of WGS in bacteriology, but it also demonstrates that concentrating WGS resources at the reference laboratory is a barrier to identifying lineages and clusters that are important at the patient care level, something that needs to be addressed in our setting in the future.

## Supporting information

Suppl Table 1 Enterobacter accession numbers

## Acknowledgements

We gratefully acknowledge clinical and diagnostic laboratory staff at the following institutions for contributing material for this study: Babcock University Teaching Hospital, Ogun State, University of Osun Teaching Hospital, Osogbo, Obafemi Awolowo University Teaching Complex, Ile-Ife, University College Hospital, Ibadan and University of Ilorin Teaching Hospital, Ilorin. We also acknowledge Tangkat Tense and Ibrahim A. Hamzat (UITH) technical contributions.

## Author Contributions

Conceptualization (AOAb, CI, DMA, INO), Data curation (FIO, AOO, OKI, VIO, IEM, AOAb, OFO, AAA, AF, BAO, CE, POO, OOO, FO, ATA, INO), Formal Analysis (FIO, AOAf, EEO), Funding acquisition (CI, DMA, INO), Investigation (FIO, AOAf, AOO, EEO), Methodology (FIO, AOAf, AOO, EEO), Project admin (AOO, AE, TO, DMA, INO), Resources (OKI, VIO, AOAb, OFO, AAA, AF, BAO, CE, POO, OOO, FO, ATA, AE, TO, CI, INO), Software (AOO, EEO, CI), Supervision (AOO, EEO, AOAb, CE, OOO, TO, CI, DMA, INO), Validation (FIO, AOO, INO), Visualization (FIO, AOAf, AOO), Writing Initial draft (FIO, INO), Editing (FIO, AOAf, BAO, OOI, AOO, SA, INO), Review of final draft (All authors).

## Ethics statement

Isolates used for this study were from positive blood cultures and collected as part of routine clinical diagnostics and surveillance in the sending laboratories of the Nigeria Antimicrobial Resistance Surveillance System from 2014 to 2020. Permission to use the isolates and their genomes for research purposes was granted by the UI/UCH ethics committee, with approval number UI/EC/22/0113.

## ACRONYMS

EHO: *E. hormaechei*
ECL: *E. cloacae*
ECA: *E. cancerogenous*
EAS: *E. asburiae*
ERO: *Enterobacter roggenkampii*
EKO: *E. kobei*
EBU: *E. bugandensis*
NR: not recorded
LUTH: Lagos University Teaching Hospital, Lagos
UCH: University college Hospital, Ibadan
OAUTHC: Obafemi Awolowo University Teaching Complex, Ile-Ife
UITH: University of Ilorin Teaching Hospital, Ilorin
UNIOSUNTH: University of Osun Teaching Hospital, Osogbo, Babcock University Teaching Hospital, Ogun State

