## Supplementary material for "Whole Genome Sequencing Reveals *Enterobacter hormaechei* as a Key Bloodstream Pathogen in Six Tertiary Care Hospitals in Southwestern Nigeria": Suppl Tables 2 to 6

**Table S2**: Initial identification of *Enterobacter hormaechei* by reference and sentinel laboratories.

| ID | Initial sentinel lab ID | VITEK2 species | % probability |
| --- | --- | --- | --- |
| UCH-OGU-P1593 | *Pseudomonas aeruginosa* | *Pseudomonas aeruginosa* | missing |
| UCH-OGU-P1573 | *Pseudomonas aeruginosa* | *Pseudomonas aeruginosa* | 98% |
| UCH-OGU-P01062 | *Enterobacteriaceae* | *Enterobacter cloacae complex* | 98% |
| UCH-OGU-P549A | *Enterobacteriaceae* | *Enterobacter cloacae complex* | missing |
| UCH-OGU-P0547A | *Enterobacteriaceae* | *Enterobacter cloacae complex* | 96% |
| UCH-OGU-P0656 | *Enterobacteriaceae* | *Enterobacter cloacae* | 95% |
| UCH-OGU-P0267C | *Enterobacteriaceae* | *Enterobacter cloacae* | 99% |
| UCH-OGU-P0144A | *Enterobacteriaceae* | *Enterobacter cloacae* | 98% |
| UCH-OGU-P1581A | *Enterobacteriaceae* | *Enterobacter cloacae* | *missing* |
| UCH-OGU-P1521 | *Klebsiella pneumoniae* | *Klebsiella pneumoniae* | *missing* |
| UCH-OGU-P1193 | *Enterobacter cloacae* | *Enterobacter cloacae* | *99%* |
| UCH-OGU-P2006 | *Enterobacter cloacae* | *Enterobacter cloacae* | *99%* |
| UCH-OGU-P1377 | *Enterobacter cloacae* | *Enterobacter cloacae* | *99%* |
| UCH-OGU-P1394 | *Enterobacter cloacae* | *Enterobacter cloacae* | *99%* |
| UCH-OGU-P1267 | *Enterobacter cloacae* | *Enterobacter cloacae* | *missing* |
| UCH-OGU-P0144B | *Acinetobacter baumannii* | *Acinetobacter baumannii* | *missing* |
| UCH-OGU-P0144C | *Acinetobacter baumannii* | *Acinetobacter baumannii* | *94%* |
| OAU-OFO-542 | *Klebsiella pneumoniae* | *Escherichia coli* | *96%* |
| OAU-AAA-578 | *Enterobacteriaceae* | *Salmonella* | *93%* |
| OAU-OFO-129A | *Enterobacteriaceae* | *Escherichia coli* | *missing* |
| OAU-OFO-575i | *Enterobacteriaceae* | *Escherichia coli* | *95%* |
| UCH-OGU-P1581B | *Enterobacteriaceae* | *Escherichia coli* | *missing* |
| LUT-BC-316 | *Staphylococcus aureus* | *Enterobacter aerogenes* | *missing* |
| OSO-OJO-S18 | *Streptococcus pyogenes* | *Escherichia coli* | *missing* |
| UCH-OGU-P1394 | *Klebsiella pneumoniae* | *Escherichia coli* | *98%* |
| UCH-OGU-P1730 | *HVN* | *Escherichia coli* | *missing* |
| LUT-BC-267 | *Acinetobacter baumannii* | *Staphylococcus haemolyticus* | *missing* |
| OSO-OJO-E2 | *Pseudomonas aeruginosa* | *Escherichia coli* | *98%* |
| UCH-OGU-P1579i | *Escherichia coli* | *Escherichia coli* | *99%* |
| UCH-OGU-19-P1403 | *Escherichia coli* | *Enterobacter cloacae* | 99% |
| UCH-OGU-19-P1400B | *Staphylococcus aureus* | *Enterobacter cloacae* | 99% |
| LUT-BC-19-196 | *Pantoea agglomerans* | *Enterobacter cloacae* | 97% |
| LUT-BC-19-506 | *Coagulase-negative staphylococcus* | *Enterobacter cloacae* | 99% |
| LUT-BC-19-584 | *Escherichia coli* | *Enterobacter cloacae* | 99% |
| OAU-OA-144 | *Enterobacter cloacae* | *Escherichia coli* | 99% |
| ELL-NOO-157B | *Klebsiella pneumoniae* | *Enterobacter cloacae* | 93% |
| ELL-NOO-172A | *Klebsiella pneumoniae* | *Enterobacter cloacae* | 94% |
| ELL-NOO-171B | *Klebsiella pneumoniae* | *Enterobacter cloacae* | 94% |
| ILO-20-ET-013 | *Escherichia coli* | *Enterobacter cloacae* | 98% |
| OAU-OA-013 | *Escherichia coli* | *Enterobacter cloacae* | 99% |
| OAU-LT-116 | *Escherichia coli* | *Enterobacter cloacae* | 99% |
| OAU-OA-020 | *Klebsiella pneumoniae* | *Enterobacter cloacae* | 98% |
| CLL-NOO-303 | *Klebsiella pneumoniae* | *Enterobacter cloacae* | 93% |
| ELL-NOO-81Bi | *Klebsiella pneumoniae* | *Enterobacter cloacae* | 98% |
| ELL-NOO-83B | *Klebsiella pneumoniae* | *Enterobacter cloacae* | 98% |
| ELL-NOO-83Bd | *Klebsiella pneumoniae* | *Enterobacter cloacae* | 99% |
| ELL-NOO-92A | *Klebsiella pneumoniae* | *Enterobacter cloacae* | 99% |
| ELL-NOO-97B | *Klebsiella pneumoniae* | *Enterobacter cloacae* | 98% |
| ELL-NOO-114B | *Klebsiella pneumoniae* | *Enterobacter cloacae* | 96% |

**Table S3**: AMR phenotypic and genotypic concordance analysis

|  | *E. hormaechei* | | | *E. cloacae* | | |
| --- | --- | --- | --- | --- | --- | --- |
| drug | concordance | specificity | sensitivity | concordance | specificity | sensitivity |
| Ampicillin | 1 | NA | 1 | 1 | NA | 1 |
| Amoxicillin/Clavulanic Acid | 0.652173913 | NA | 0.652174 | 0.933333 | 0 | 1 |
| Piperacillin/Tazobactam | 0.538461538 | 0.416667 | 0.642857 | 0.4 | 0 | 1 |
| Cefuroxime | 1 | NA | 1 | 0.733333 | NA | 0.733333 |
| Cefuroxime Axetil | 1 | NA | 1 | 0.733333 | NA | 0.733333 |
| Ceftriaxone | 0.72 | 0 | 1 | 0.866667 | 0.75 | 0.909091 |
| Cefoperazone/Sulbactam | 0.307692308 | 0 | 1 | 0.266667 | 0.266667 | NA |
| Cefepime | 0.423076923 | 0 | 1 | 0.733333 | 0.5 | 0.888889 |
| Ertapenem | 0.869565217 | 1 | 0.4 | - | - | - |
| Imipenem | 1 | 1 | 1 | - | - | - |
| Meropenem | 1 | 1 | 1 | - | - | - |
| Amikacin | 1 | 1 | 1 | 0.333333 | 0.333333 | NA |
| Gentamicin | 0.923076923 | 0.888889 | 0.941176 | 1 | 1 | 1 |
| Colistin | 0.75 | 1 | 0 | 0 | NA | 0 |
| Trimethoprim/Sulfamethoxazole | 0.833333333 | 0.6 | 1 | 1 | 1 | 1 |

**Table S4**: Numbers of Enterobacter genomes and AmpC variants (using AMRFinderPlus and CARD)

| *Enterobacter* spp. | No. Of genomes | No. of AmpC variants | Assigned ACT, CMH, and MIR variant(s) |
| --- | --- | --- | --- |
| *E. hormaechei* | 61 | 14 | ACT-15(4), -16(8), -17(2), -24(4), -25(11), -41(1), -45(6), -46(4), -61(2)-69(2) -70(1), -74(7) -75(5), -84(4) |
| *E. cloacae* | 26 | 3 | CMH-1(1), -3(7), -4(15), -7(3) |
| *E. roggenkampii* | 4 | 3 | MIR-2(2), -23(1), ACT-62(1), |
| *E. bugandensis* | 3 | 3 | ACT-72(1), -77(1), -78(1) |
| *E. kobei* | 2 | 2 | ACT-28(1), ACT-9(1) |
| *E. asburiae* | 1 | 1 | ACT-2(1) |
| *E. cancerogenous* | 1 | 1 | ACT-8(1) |

*The number of strains with each variant is in brackets

**Table S5**: AST data of outbreak strains

| **ST109-NDM** | **Alternative ID** | **Amoxicillin/Clavulanic Acid** | **Piperacillin/Tazobactam** | **Cefuroxime** | **Cefuroxime Axetil** | **Ceftriaxone** | **Cefoperazone/Sulbactam** | **Cefepime** | **Ertapenem** | **Imipenem** | **Meropenem** | **Amikacin** | **Gentamicin** | **Nalidixic Acid** | **Ciprofloxacin** | **Tigecycline** | **Nitrofurantoin** | **Trimethoprim/Sulfamethoxaz** |
| --- | --- | --- | --- | --- | --- | --- | --- | --- | --- | --- | --- | --- | --- | --- | --- | --- | --- | --- |
| **UCH-OGU-P0144A** | **G18503214** | **>= 32/R** | **>= 128/R** | **>= 64/R** | **>= 64/R** | **>= 64/R** | **>= 64/R** | **16/R** | **>= 8/R** | **>= 16/R** | **>= 16/R** | **>= 64/R** | **>= 16/R** | **>= 32/R** | **<= 0.25/R** | **<= 0.5/R** | **64/I** | **<= 20/S** |
| **UCH-OGU-P0656** | **G18503210** | **>= 32/R** | **>= 128/R** | **>= 64/R** | **>= 64/R** | **>= 64/R** | **>= 64/R** | **8/R** | **>= 8/R** | **>= 16/R** | **>= 16/R** | **>= 64/R** | **>= 16/R** | **>= 32/R** | **<= 0.25/S** | **1/S** | **64/I** | **<= 20/S** |
| **UCH-OGU-P0144C** | **G18581058** |  | **>= 128/R** |  |  | **>= 64/R** | **>= 64/R** | **16/I** |  | **>= 16/R** | **>= 16/R** |  | **>= 16/R** |  | **<= 0.25/R** | **<= 0.5/S** |  | **<= 20/S** |
| **ST148** |  |  |  |  |  |  |  |  |  |  |  |  |  |  |  |  |  |  |
| **UCH-OGU-P01062** | **G18503206** | **>= 32/R** | **16/S** | **>= 64/R** | **>= 64/R** | **>= 64/R** | **<= 8/S** | **4/S** | **<= 0.5/S** | **1/S** | **<= 0.25/S** | **<= 2/S** | **>= 16/R** | **16/S** | **1/S** | **2/S** | **64/I** | **>= 320/R** |
| **UCH-OGU-P1581B** | **G18593207** | **>= 32/R** | **32/I** | **>= 64/R** | **>= 64/R** | **>= 64/R** | **<= 8/S** | **2/S** | **<= 0.5/S** | **1/S** | **<= 0.25/S** | **<= 2/S** | **>= 16/R** | **16/S** | **1/S** | **2/S** | **128/R** | **>= 320/R** |
| **UCH-OGU-P0267C** | **G18503213** | **>= 32/R** | **32/I** | **>= 64/R** | **>= 64/R** | **>= 64/R** | **<= 8/S** | **2/S** | **<= 0.5/S** | **0.5/S** | **<= 0.25/S** | **<= 2/S** | **>= 16/R** | **16/S** | **1/S** | **2/S** | **64/I** | **>= 320/R** |
| **UCH-OGU-P1573** | **G18501064** |  | **32/S** |  |  |  | **<= 8/S** | **32/R** |  | **0.5/S** | **<= 0.25/S** | **4/S** | **>= 16/R** |  | **1/S** | **1/R** |  |  |
| **UCH-OGU-P1593** | **G18501062** |  | **<= 4/S** |  |  |  | **<= 8/S** | **<= 1/S** |  | **1/S** | **<= 0.25/S** | **<= 2/S** | **<= 1/S** |  | **<= 0.25/S** | **1/R** |  |  |

**Table S6**: Vitek2 test specificity, sensitivity, positive predictive values, and negative predictive values

|  | *E. hormaechei* | *E. cloacae* |
| --- | --- | --- |
| % sensitivity | 0.0 | 1 |
| % specificity | 0.0 | 0.0 |
| % positive predictive value | 0.0 | 1 |
| % negative predictive value | 0.0 | 0.0 |
