## Supplementary material for "Whole Genome Sequencing Reveals *Enterobacter hormaechei* as a Key Bloodstream Pathogen in Six Tertiary Care Hospitals in Southwestern Nigeria": Suppl Figure 1

| ST109-ndm | G20500059 | G18581057 | G18503214 | G18503210 | G18581058 | G18503419 |
| --- | --- | --- | --- | --- | --- | --- |
| G20500016 | 0 | 27725 | 27703 | 27748 | 27676 | 31068 |
| G20500059 | 0 | 27578 | 27561 | 27608 | 27542 | 31035 |
| G18581057 | 27578 | 0 | 0 | 0 | 1 | 31383 |
| G18503214 | 27561 | 0 | 0 | 0 | 1 | 31396 |
| G18503210 | 27608 | 0 | 0 | 0 | 1 | 31432 |
| G18581058 | 27542 | 1 | 1 | 1 | 0 | 31376 |
| G18503419 | 31035 | 31383 | 31396 | 31432 | 31376 | 0 |
| G20500210 | 31161 | 31461 | 31477 | 31513 | 31452 | 0 |

| ST148 | G18503206 | G18593207 | G18503213 | G18501064 | G18501062 | G20501659 | G18503413 |
| --- | --- | --- | --- | --- | --- | --- | --- |
| G18503206 | 0 | 0 | 0 | 0 | 1 | 174 | 143 |
| G18593207 | 0 | 0 | 0 | 0 | 1 | 170 | 144 |
| G18503213 | 0 | 0 | 0 | 0 | 1 | 170 | 144 |
| G18501064 | 0 | 0 | 0 | 0 | 1 | 170 | 144 |
| G18501062 | 1 | 1 | 1 | 1 | 0 | 164 | 145 |
| G20501659 | 174 | 170 | 170 | 170 | 164 | 0 | 254 |
| G18503413 | 143 | 144 | 144 | 144 | 145 | 254 | 0 |

**Figure S1**: SNP distances for ST109 and ST148 strains with likely outbreak strains in darker shade
